# Lévy movements and a slowly decaying memory allow efficient collective learning in groups of interacting foragers

**DOI:** 10.1101/2023.05.08.539904

**Authors:** Andrea Falcón-Cortés, Denis Boyer, Maximino Aldana, Gabriel Ramos-Fernández

## Abstract

Many animal species benefit from spatial learning to adapt their foraging movements to the distribution of resources. Learning involves the collection, storage and retrieval of information, and depends on both the random search strategies employed and the memory capacities of the individual. For animals living in social groups, spatial learning can be further enhanced by information transfer among group members. However, how individual behavior affects the emergence of collective states of learning is still poorly understood. Here, with the help of a spatially explicit agent-based model where individuals transfer information to their peers, we analyze the effects on the use of resources of varying memory capacities in combination with different exploration strategies, such as ordinary random walks and Lévy flights. We find that individual Lévy displacements associated with a slow memory decay lead to a very rapid collective response, a high group cohesion and to an optimal exploitation of the best resource patches in static but complex environments, even when the interaction rate among individuals is low.

**Author Summary:** How groups of social animals collectively learn to find and exploit resources in complex environments is not well-understood. By means of a computational model where individuals are initially spread out across a landscape, we study the effects of individual exploratory behaviors and memory capacities on the emergence of spatial learning. Collective learning emerges spontaneously only if group members transfer information between each other at a sufficiently high rate, so that individual experiences can be used by others. In static but heterogeneous environments with many resource sites of varying attractiveness, we find that random displacements over many spatial scales combined with a slow memory decay lead to a rapid collective response and highly cohesive groups. Collective learning is noticeable through an optimal exploitation of the best resource sites, which far exceeds what individuals would achieve on their own. Our study sheds light on important mechanisms responsible for collective learning in ecology, with potential applications in other areas of science.

## INTRODUCTION

Learning is evidenced through changes in animal behaviour as a result of experience and is well documented in many species. Learning brings numerous ecological advantages by improving the foraging efficiency [1], increasing the ability to escape from predators [2] or through the choice of major low-cost migrations corridors between seasonal ranges [3, 4], for instance. Learning processes are comprised of several steps that are necessary for a proper information acquisition based on experience, such as the collection, storage and retrieval of information. These steps are directly conditioned by the intrinsic cognitive capacities of the individual, and, in the case of foraging, the strategies used to explore the environment [5]. A learning process is deemed successful when the organism has the ability to choose among various options with a bias toward the most rewarding ones, based on previous experience. During the process, past events can be recent or remembered over long periods of time [6, 7]. In this sense, we can define learning as in the context of machine learning: improved performance on a specific task due to prior experience [5].

The learning of spatial features in an environment is closely related to exploratory behavior. Some strategies may be more beneficial than others to collect information across different spatial and temporal scales. A body of studies have reported Lévy patterns in the foraging movements of diverse animal species [8–13]. Lévy flights (LF) are random sequences of independent displacements whose lengths are drawn from a probability distribution function having a power-law tail [14, 15]. An important statistical feature of LF is scale invariance in space and time [16], which makes them a convenient modelling tool to analyze animal paths that involve multiple spatial or temporal scales. Such trajectories quickly diffuse in space and can be depicted as composed of many random walks with small, local displacements connected to each other by displacements of widely varying lengths [11]. Although there is controversy about the validity of the Lévy-flight approximation to describe the motion of animal groups [17–19], there is substantial evidence that the Lévy-flight approximation works well for some species. Indeed, patterns that can be accurately described by means of Lévy flights (or related Lévy walks) have been observed in the movements of seabirds [8, 20], marine predators [21], foraging bumble bees [9], African jackals [22], spider monkeys [12], or Drosophila flying in a small circular arena [23], among others. It is not our purpose in this article to delve into the debate about the validity of the Lévy-flight approximation. Instead, we will take for granted that Lévy flights do exist as a model of multiple-scale animal movement, and will explore the consequences of this hypothesis regarding foraging with memory and learning.

As mentioned above, important steps during a learning process are the storage and retrieval of information, two cognitive processes that depend on the memory capacity of the organism. Spatial memory is documented in many animal species, from large herbivores to birds [1, 24–26]. The use of memory, as well as its decay over time, can be inferred by fitting models to real trajectories obtained from tracking devices in foraging landscapes where the resource patches are well identified [27, 28]. Recent theoretical approaches incorporating memory-based movements into ordinaryrandom-walk models have shown how spatial learning can emerge in principle [29–31]. This phenomenon is noticeable by frequent revisits to certain places where resources are located [32] or through the emergence of home ranges and preferred travel routes [33, 34].

Theoretical studies often assume that individuals have an infinite memory capacity, *i*.*e*., the ability to remember all the sites they have visited throughout their lifetimes. Such simplification has allowed a better understanding of the effects of memory on space use and the emergence of site fidelity. However, it is well known that memory actually decays in time: it is easier to remember recent events than those that happened far in the past. Memory decay has been incorporated into optimal foraging theory [35], and it has been well documented empirically in many animal species such as coal tits [36], desert ants [37], golden lion tamarins and Wied’s marmosets [38]. In the case of bison [1], a memory kernel slowly decaying with time as a power-law better described the revisitation patterns than the more standard exponential kernel. Memory decay is thought to be advantageous when the environment is changing over time [35] and can enhance the foraging success of primates because it allows animals to constantly update the relevant features of their environment [38].

The studies described above have mainly focused on individual learning. However, many animal species live in groups and sociability brings many benefits, ranging from a reduced predation risk [39–42] to an increased foraging efficiency in uncertain environments [43–46]. The concept of collective learning describes any learning process that is mediated by the transfer of information, skills or strategies that were obtained in social contexts. Such interactions can modify individual behavior and lead to coordinated collective dynamics [5]. There are several types of collective learning, but all of them include the exchange of useful information in a direct or indirect way. Individuals can increase their probability of adopting a certain behavior in the presence of a peer, like in the case of bison, which are more likely to travel to a given new location when another animal in the group knows about this location. [47]. The interest of an individual about a particular location can arouse the interest of others, as in elephants, where matriarchs lead herds to waterholes unknown by the rest of the group [48], or in killer whales, where old females tend to lead the displacements of the group when prey abundance is low [49]. Novel individuals can copy a model behavior through observation that results in a reliable and similar outcome, as in fish where naive individuals learn migration routes through associations with experienced individuals [50]. Given that individual and collective learning could benefit from each other [51–54], we extend the definition of learning exposed above to define collective learning as: the improved performance of a group to solve a specific task due to the accumulation of individual experience and the transfer of social knowledge.

Several spatially explicit models have shown that spatial learning in groups can be affected by the rates at which individuals interact or use previous information [52–54]. For instance, collectives of interacting random walkers with infinite memory capacities were studied in [52]. In heterogeneous environments, these models naturally exhibit collective learning over a wide range of parameters, a phenomenon noticeable by the capacity of the foraging group as a whole to strongly aggregate and localize around the richest resource sites available in the environment.

Despite these advances, there is still a lack of understanding on how different combinations of individual exploration types and memory capacities affect the emergence of collective learning. Here, we computationally analyze a model swarm without a leader, whose task is to find and exploit the resource sites of highest reward in an environment containing many resource sites. In particular, we study the effects of forager individual dispersion, modelled by LF movements of varying exponent, combined with a power-law memory decay, on the emergence of collective learning through information transfer, and contrast the results with those obtained from movements of constant step length or perfect memory [52]. We find that LF dynamics and a moderate memory decay allow for a fast and effective collective learning in complex environments, even if those are static.

## MODEL

The collective foraging model presented here extends the one studied in [52]. Briefly, the motion of individuals is assumed to be driven by the combination of two basic movement modes: standard random walk displacements (with probability 1 *− q*) and preferential returns to places visited in the past (with probability *q*). In the latter mode, an individual chooses a particular site from its own experience (with probability 1 − *ρ*) or from the experience of another group member (with probability *ρ*). The probability of choosing a particular site for revisiting is proportional to the accumulated amount of time that the individual (or its peer) has spent at that site. Therefore, the sites that are often occupied have a higher probability of being revisited in the future: they are linearly reinforced [32, 52]. Since this preferential dynamics depends on the whole history, it implies that each individual in the swarm can remember which sites were previously visited as well as the duration of these visits, either by its own experience (memory) or by *communication* with peers (social interaction). Such communication takes place on an information transfer network in which the agents are represented by nodes, and the links between agents allow information exchange.

### Environment

We consider a discrete lattice of *L × L* sites in two dimensions, with unit spacing and reflective boundaries. The coordinates of a site *n* are a pair of integers (*x, y*). On the lattice, resources (or targets) are randomly distributed with density *δ <* 1, hence, on average, there are *M* = *δL*^2^ targets in the system. To each target *i* we assign a fixed weight or attractiveness *γ*_*i*_ (related to the site “reward” in the context of learning), which is a random number uniformly distributed in the interval (0, *γ*_*max*_), where *γ*_*max*_ *<* 1 is a given parameter. Let 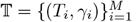 denote the set that contains the positions *T*_*i*_ and weights *γ*_*i*_ of the targets. The target of highest weight, *γ*_best_ = max_*i*=1,…,*M*_ *{γ*_*i*_*}*, is the best target and is denoted as T_best_. Although the targets are spatially distributed randomly across the environment, heterogeneity is introduced through the attractiveness parameter *γ*_*i*_. Then, different targets can distinctly contribute to the system’s dynamics. Other choices for the probability function *P* (*γ*) are possible, such as a power-law probability distribution *P* (*γ*) = *Cγ*^*− α*^. In this case, there would be a few highly attractive targets, while most are poorly attractive. Previous work shows that food distribution in certain environments can be approximated by a power-law [55]. However, this work presents results for a uniform probability distribution *P* (*γ*), which also includes a certain amount of heterogeneity.

### Foragers

We consider *N* walkers with random initial positions on the lattice, connected by a complete communication network. Namely, every walker can communicate with any other group member [**?**]. The time variable *t* is discrete and during a time step *t → t* + 1, each walker *l* = 1, …, *N* updates its position 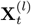 as follows.

i. *Self-memory mode*: If not on a target, with probability *q*(1 *− ρ*) the walker moves to a previously visited site, that is: 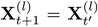 where *t*′ is an integer in the interval [0, *t*] chosen according to a probability distribution *p*_*t*_(*t*′) given *a priori* and denoted as:

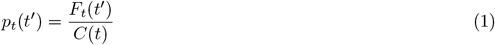

with 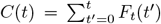 a normalization factor and *F*_*t*_ (*t*′) a memory kernel function. Here we consider a power-law memory decay given by

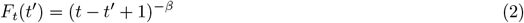

where *β* ≥ 0. If *β* = 0, one recovers the linear preferential visit model with non-decaying memory [52], where the probability to visit a site is simply proportional to the total amount of time spent on that site up to *t*. With *β >* 0, it is more likely that the individual chooses a recently visited place for relocation, rather than a site visited long ago. The rule in Eq. (2) is then equivalent to assuming that each site (target or not) has an accumulated weight for the forager, such that each visit to a given site (at time *t*′) increments its weight in 1. At later times (*t > t*′), this increment decreases from its initial value 1 and tends to zero as a power-law. Hence, frequently visited sites have a larger accumulated weight and are more likely to be revisited again (preferential visit mechanism) but sites that have not been visited for a long time have vanishing weights and tend to be forgotten by the forager. In addition to the power-law memory kernel above, we also performed simulations with an exponential kernel 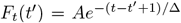 with no qualitative difference in the results (see Supporting Information, Fig A in S1 Text).
ii. *Information transfer mode*: If not on a target, with probability *qρ* the walker *l* randomly chooses another walker (*m*) and relocates to a place already visited by *m*, according to the same preferential rule with memory decay: 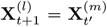 where *t*′ is again a random integer in the interval [0, *t*] chosen from the probability distribution *p*_*t*_(*t*′)
iii. *Random motion mode*: If not on a target, with probability 1 *− q* the walker *l* performs a random displacement, or 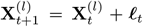. Here, ***ℓ***_*t*_ = (*ℓ*_*x*_, *ℓ*_*y*_), where *ℓ*_*x*_ and *ℓ*_*y*_ are two integers independently drawn from the distribution:

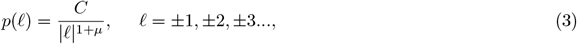

with *C* a normalization constant and 0 ≤*µ*≤ 2. When *µ*≥ 2, the random walk process becomes asymptotically Gaussian, similarly to an ordinary random walk with small jumps, *e*.*g*., to nearest neighbors (n.n.) lattice sites. Conversely, when *µ*→ 0, the dynamics approximately correspond to random relocation in space, where very large displacements across the landscape are frequent. For intermediate values of *µ*, self-similar Lévy flights are generated. This random movement mode does not take into account travel costs. In the Supplementary Information, we present a variant of this rule that introduces cost, thus discouraging long movement steps (see Fig B in S1 Text and the Discussion below). The main results with and without travel costs are qualitatively the same. Therefore, in all the results presented in what follows, we do not take into account any kind of cost function for the motion of the particles.
iv. Feeding: If on a target (*i*.*e*. if there exists an *i* such that 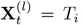), the walker *l* stays on that site with probability *γ*_*i*_, and with the complementary probability 1 *− γ*_*i*_ leaves the site using one of the movement rules (*i*) *−* (*iii*) above with their respective probabilities.

In summary, the parameter *q* represents the probability of memory use (while 1*− q* is the probability to perform a random step), whereas *ρ* is the interaction parameter, or the probability that the walker uses, in the memory mode, the experience of another individual. The mean time spent feeding on a target *T*_*i*_ during a visit is 1*/*(1 *− γ*_*i*_), which can be considered as the mean reward associated to *T*_*i*_. Hence a site with a *γ*_*i*_ close to *γ*_*max*_ (which slightly below 1) represents a “rich” site. By setting *β* = 0 (perfect memory) and if the jumps in the random motion mode are limited to nearest neighbours (n.n.) lattice sites, one recovers the model of ref. [52]. See also refs. [53, 54] for other variants of this model.

## RESULTS

With the simple rules described above, collective learning may emerge via the localization of a large fraction of the individuals around a good resource site, chosen among the many resource sites available in the foraging domain (phenomenon known as *selective localization*). Fig 1 displays an example of spatial configuration obtained at large time in a simulation by iterating the rules (i)–(iv), for a swarm of interacting foragers with non-decaying memory and performing Lévy flights with index *µ* = 0.5. One notices a highly non-uniform distribution of foragers around a few resource sites of high attractiveness.

**Fig 1.**
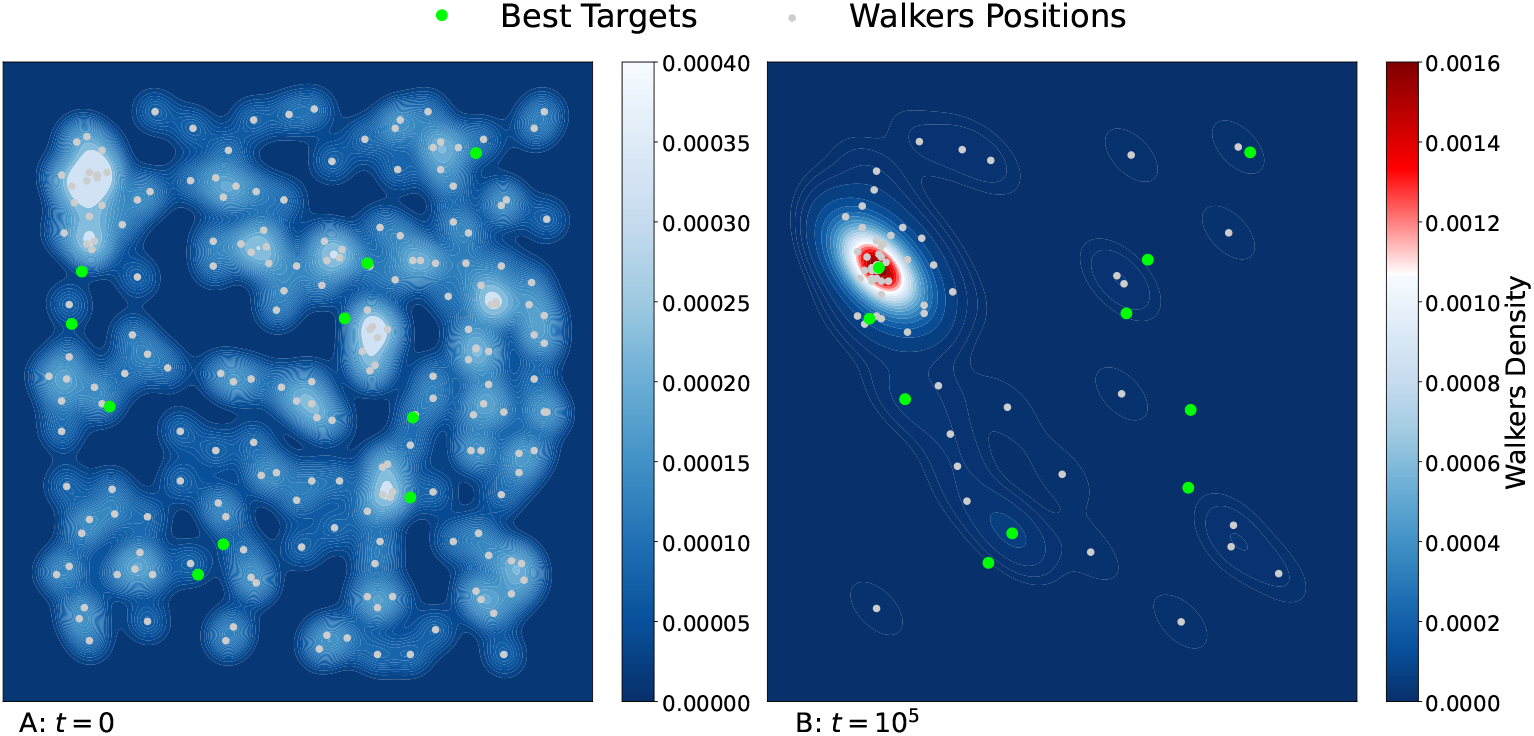
Initial (A) and final (B) positions of 200 walkers (white dots) following the dynamical rules (i)–(iv), after *t* = 10^5^ iterations in a two-dimensional environment of size *L* = 200. The most attractive resource sites (with *γ ≥* 0.85) are represented by green circles (among a total of 100). The values of the other parameters are *q* = 0.25, *β* = 0, *ρ* = 0.5, *γ*_*max*_ = 0.9, and *µ* = 0.5. Initially, all the walkers are randomly distributed across the square lattice. Note that at the end of the simulation, most of the walkers aggregate around a very attractive resource site, as indicated by the color gradients showing the density of walkers.

### Infinite Memory (*β* = 0)

Fig 2A and 2B show the occupation probability 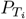 of each target *T*_*i*_ as a function of its weight *γ*_*i*_, for nearest neighbour (n.n) and LF random jumps, respectively. At large times, a steady state is reached and 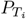 is defined as the probability that a walker chosen at random occupies the target *T*_*i*_. This quantity gives us a measure of how selectively foragers choose their resources. When interactions and memory are at play, 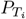 acquires its largest value for *γ*_*i*_ = *γ*_max_ and decays very quickly as *γ*_*i*_ decreases. In the n.n. case (Fig 2A), the occupation probability of the resource site T_best_, denoted as 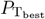, rises to about 0.25 in certain environments. Therefore, there is a probability of order 1 to find any forager (at a given time) on the site giving the maximum reward, out of the *L*^2^ = 40, 000 visitable sites. For the LF dynamics (Fig 2B), 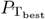 grows to about 0.35 in some environments. It is important to notice that when the swarm individuals do not communicate with each other (pink squares), one still observes some localization around the best resource site with LF jumps, whereas this phenomenon almost vanishes with n.n. jumps.

**Fig 2.**
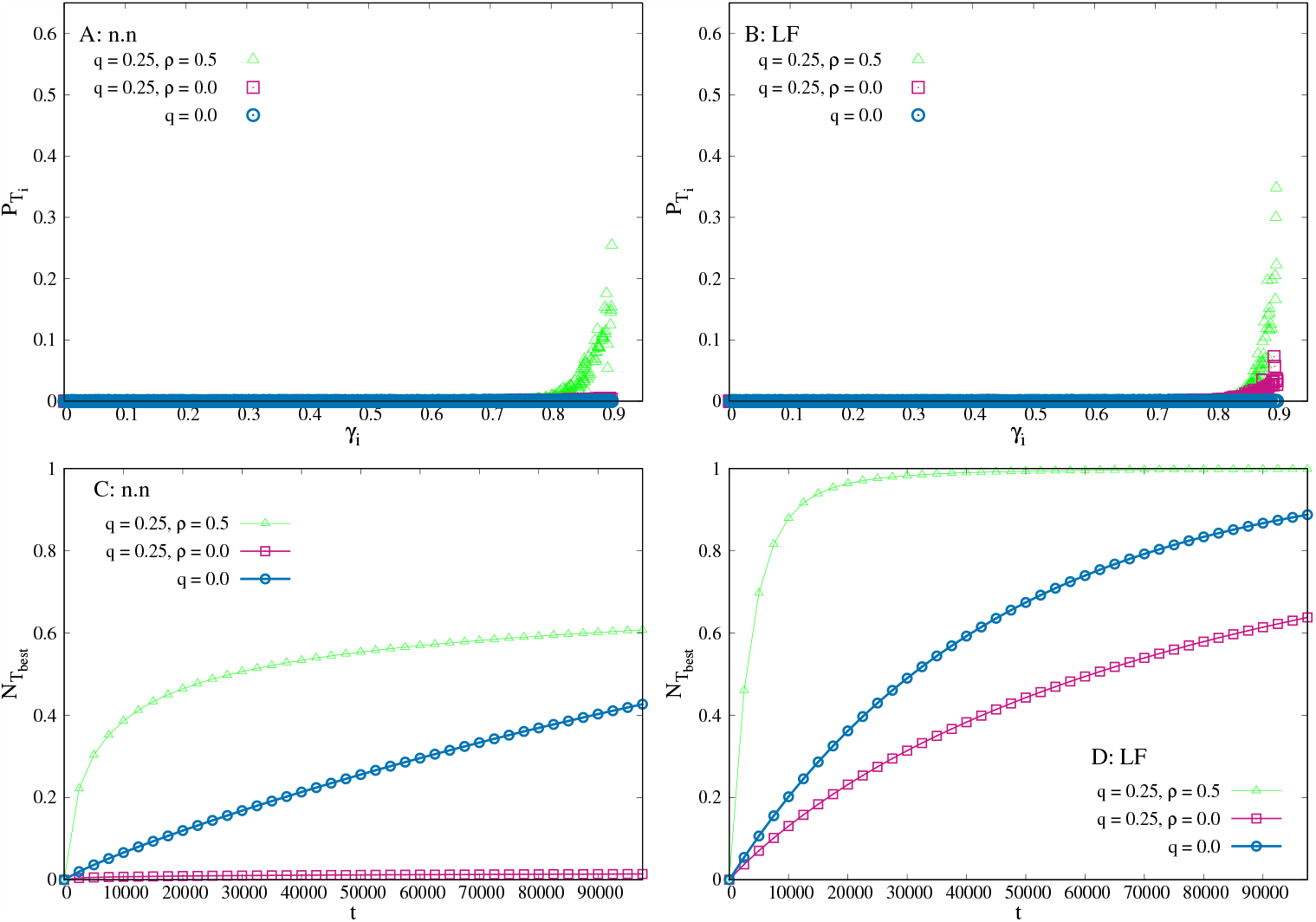
Infinite memory case. Simulations with *N* = 200 walkers in an environment with 200 *×* 200 lattice sites, *δ* = 0.0025 (*M* = 100 resource sites), *γ*_*max*_ = 0.9 and *t* = 10^5^. A) Final target occupation probability 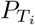 for n.n. dynamics vs. the target weight *γ*_*i*_. Each point represents a target. B) Same quantity as in (a) for LF dynamics with index *µ* = 0.5. C) Fraction of swarm individuals that have visited the best target site at least once at time *t*, for different memory and communication rates. D) Same quantity as in C) for Lévy flight dynamics with index *µ* = 0.5. 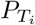 and 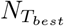 (*t*) are obtained from averaging over 1000 independent dynamics in a same environment. The data from 10 different environments are aggregated in A) and B), and averaged in C) and D). In each panel, the green triangles correspond to a system in which there is communication between particles (*ρ* = 0.5) and memory (*q* = 0.25); pink squares correspond to memory (*q* = 0.25) but no communication (*ρ* = 0); and the blue circles correspond to neither memory nor communication (random walks of independent particles, *q* = 0, *ρ* = 0).

When the foragers do not use memory (*q* = 0, blue circles in Fig 2 for n.n. and LF), the selectivity disappears altogether. In this case, the foragers are independent and memory-less random walkers. Their occupation probability of a site *i* in the steady state can be calculated exactly and is given by 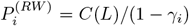, where *γ*_*i*_ = 0 for a non-target site, where 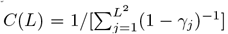 is a normalization factor. This result is independent of the step distribution *p*(*ℓ*). As *C*(*L*) *∝* 1*/L*^2^, the occupation probability of any site is very small, even for *γ*_*i*_ = *γ*_*max*_. One can also show that if these walkers interact (*e*.*g*., every individual can jump with probability *ρ* to the site currently occupied by another forager chosen at random), the asymptotic occupation probability is given by the same 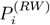 as above. Therefore, when there is no memory, communication between particles does no contribute at all to the dynamics of the system. The emergence of selective localization in our model with n.n. dynamics is thus only possible with both memory use and communication, while for the LF dynamics, individual memory use is sufficient.

Fig 2C and 2D display the fraction of individuals (denoted as 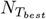 (*t*)) that have visited the best site at least once at time *t*, for n.n. and LF swarms, respectively. For a n.n. swarm with memory but without communication (*q* = 0.25 and *ρ* = 0, pink squares), very few individuals find the best target site, even after a long period of time. This is because with small steps, memory causes each individual to explore intensively a limited region of space around its starting point, which is unlikely to contain the best site. In the absence of memory and communication, the agents perform pure random walks (*q* = 0, blue circles) and can visit the best target just by chance. Under such conditions, 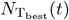 grows linearly with *t* at short times. However, in the same swarm with memory and information transfer (*q* = 0.25 and *ρ* = 0.5, green triangles), a much larger number of individuals rapidly find the best target, although this fraction remains well below 100% in all the configurations that we explored. In Fig 2D, LF dynamics exhibit similar trends. In this case, though, even in the absence of communication many individuals that use memory can find the best target. Thanks to LFs, individuals actually explore distant parts of the environment and can reach the best target by themselves more easily. Importantly, in LF swarms with memory and communication, after a short time, almost the 100% of the individuals have encountered the best target.

The quantity 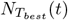 (*t*) also gives access to a dynamical quantity of particular interest in random search processes, namely, the mean time needed by an individual to visit a given site (here, the best resource site) for the first time. In Fig 2C and 2D, this mean first passage time is actually equal to the area (integrated over the time interval [0, *∞*)) between the curve of the fraction of individuals 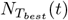 (*t*) and the horizontal line of coordinate unity. This is a very general equality, valid for any stochastic process [56]. Clearly, the first passage time is considerably shorter for LF than for n.n. swarms, where it can be huge due to the fact that 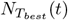 (*t*) very slowly approaches 1. Among LFs, the shortest first passage times are observed with both memory and interactions.

#### Behavior as a function of the information transfer rate ρ

Fig 3A displays the variations with *ρ* of the occupation probability 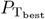 for both n.n. (pink) and LF (blue) dynamics. This probability grows rapidly with *ρ* and reaches a plateau, whose value is higher for LF than for n.n. walkers. Note also that, to achieve the plateau value, LF swarms need a larger communication rate (*ρ ≥* 0.2) than n.n. walkers (*ρ ≥* 0.1).

**Fig 3.**
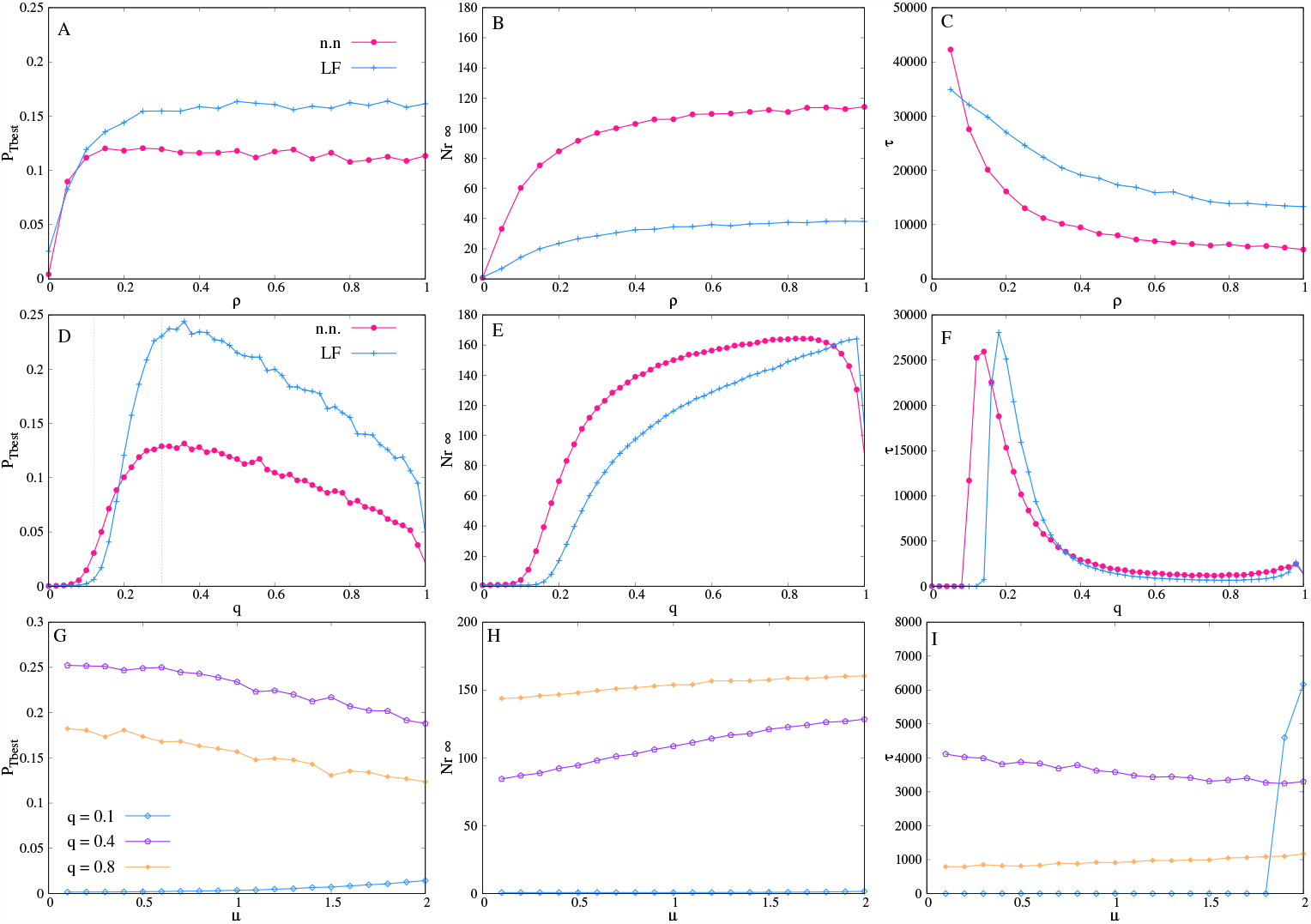
Infinite memory case. Occupation probability of the best target, 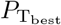 (left column), asymptotic cohesion 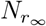 (center column) and learning time *τ* (right column), as function of: (A,B,C) the information transfer rate *ρ*, with *q* = 0.25 and *µ* = 0.5; (D,E,F) the rate of memory use *q*, with *ρ* = 0.5 and *µ* = 0.5; and G-I the Lévy index *µ*, for different values of memory use. In panel D, the values of the threshold probability *q*_*t*_ (green dashed line) and the optimal probability *q*^*∗*^ (blue dashed line) are indicated for the curve corresponding to Lévy flights swarms.The other parameters are those of Fig 2. All the curves are averages over ten different landscape configurations.

Fig 3B shows the asymptotic cohesion reached by the swarm at long times, denoted as 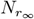, as a function of the interaction parameter *ρ*. To compute 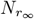, we define the neighborhood of a forager by a disk of radius 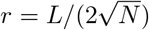 centered at the forager position. The length *r* represents half of the mean distance between two neighboring foragers if they were distributed randomly in space with homogeneous probability. For each forager, at time *t*, we determine the number of other group members located in its neighborhood and take the average over all foragers, denoted as *N*_*r*_(*t*) [52]. We define 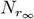 as the limit of *N*_*r*_(*t*) when *t → ∞*, in the steady state. As can be observed in Fig 3B, this quantity also reaches a plateau as *ρ* increases. The swarms with n.n. dynamics are very cohesive. This is expected because in the n.n. dynamics, the frequent use of memory (*q* = 0.25) does not give the time for the individuals to diffuse away from the best resource sites. On the other hand, the large jumps of the Lévy processes are likely to spread the group further around these sites, producing less cohesion. Nevertheless, LF swarms remain more cohesive than foragers randomly distributed in space, as exemplified by Fig 1.

Fig 3C displays the typical time taken for the swarm to reach half of its asymptotic cohesion. We denote this quantity as *τ*, which is also a measure of the time taken by the collective learning process, *i*.*e*., the adaption time of the group to the environment. The variation of *τ* as a function of *ρ* exhibit a slow decay until a finite asymptotic value is reached. Hence, more frequent interactions accelerate the convergence toward collective learning, which is faster for n.n. groups than for LF groups.

#### Behavior as a function of the memory use rate q

In panels D to F of Fig 3, we fix *ρ* = 0.5 and study the influence of *q* (the ratio of memory use) on the quantities 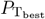, 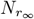 and *τ*. Fig 3D displays the occupation probability 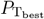 as a function of *q*. There exists a threshold value *q*_*t*_ above which the swarm presents collective learning, and this value is slightly lower for n.n. (*q*_*t*_ *≈* 0.08) than for LF dynamics (*q*_*t*_ *≈* 0.12). After the learning threshold, 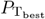 reaches an optimum at a certain *q* = *q*^*∗*^. For *q ≳* 0.2, LF swarms exhibit a better performance than n.n. swarms. In both cases, an excessive use of memory leads to a degraded localization around the best target site. This is because when *q →* 1, each individual tends to get “trapped” in the first visited site, reducing its exploration capacity to zero. Fig 3E shows that, as in Fig 3B, the cohesion of the n.n. swarms is larger than for LF swarms. In both cases, the asymptotic cohesion of the swarm decays abruptly when *q* approaches unity. In Fig 3F, the learning time *τ* exhibits a maximum immediately after the learning threshold, whereas it further decays until reaching a much lower asymptotic value, that is roughly the same in both dynamics.

#### Behavior as a function of the Lévy flight index µ (length of the steps)

These results are further complemented by studying the effect of the Lévy index *µ* on the dynamics of the system, as shown in Fig 3G, H and I. At medium rates of memory use (*e*.*g*., purple line of Fig 3G), localization improves as the flights become super-diffusive (smaller *µ*), despite of being less cohesive (Fig 3H) and slightly slower in learning (Fig 3I). Thus, the exploratory flights with shorter step lengths (larger *µ*), and hence closer to n.n. dynamics, are slightly more cohesive, and learn slightly faster. At small and large rates of memory use, localization and group cohesion are very weak, while *τ* is small because no, or little, aggregation occurs.

### Memory decay (*β >* 0)

The occupation probability of each target as a function of its attractiveness *γ*_*i*_ is now shown in Fig 4 for a non-zero value of the memory decay parameter (*β* = 1, see Eq. 2). In Fig 4A and 4B we show 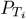 for n.n. and LF swarms, respectively. As in the perfect memory case, 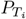 is very small for almost all *γ*_*i*_’s but becomes very large (of order 1) when *γ*_*i*_ is close to *γ*_*max*_, provided that memory use and communication are at play in the group (grey triangles). Again, our model foragers are attracted to the best target sites and localize very selectively. When communication is suppressed (*ρ* = 0, blue circles) 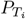 drops abruptly for all *γ*_*i*_ in n.n swarms. For LF swarms, though, the lack of communication does not destroy selectivity target localization, but decreases it by a factor of about 3 (see also Fig 5A, green curve). When the individuals are memory-less, (*q* = 0, green crosses), selectivity completely disappears in both dynamics, as expected. By comparing with Fig 2, one notices that memory decay in the LF swarm markedly improves the selective localization on good targets compared to the case of infinite memory (see a value of 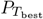 above 0.60 in Fig 4B). Although memory is crucial, allowing the individuals to forget about those sites visited in the far past, gives the group opportunities to find and exploit better resource sites, even in a static environment.

**Fig 4.**
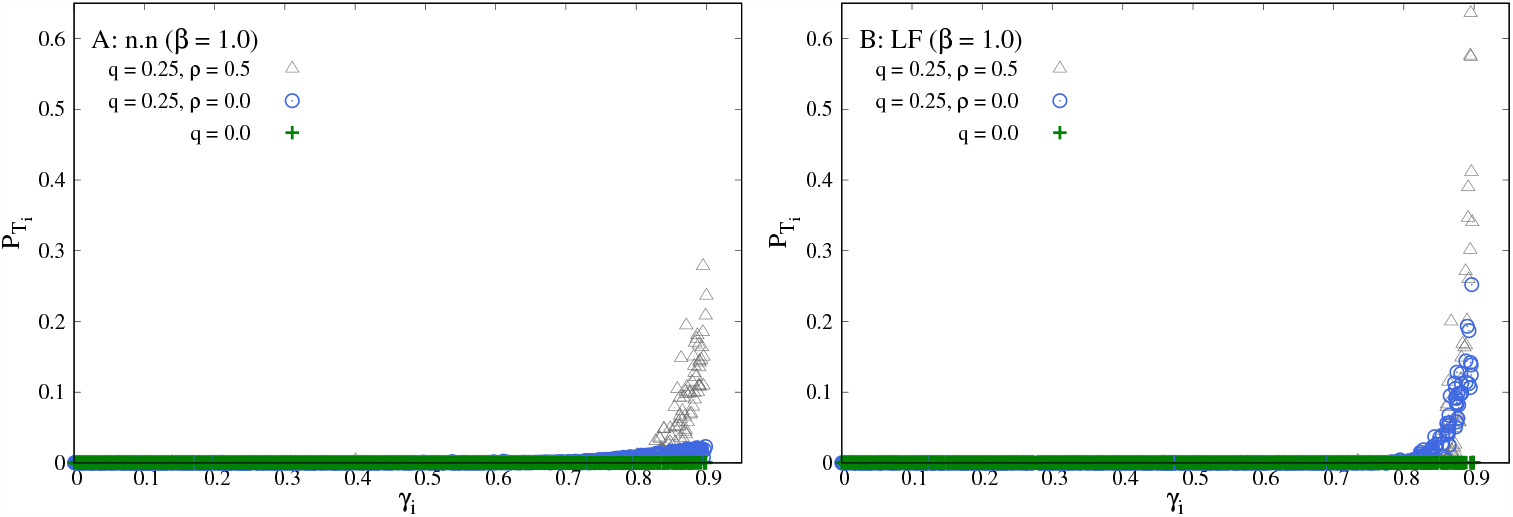
Memory decay case. A) Final target occupation probability 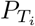 for n.n. dynamics vs. the value *γ*_*i*_ of the target attractiveness. B) Same as A) but for the Lévy flight dynamic with *µ* = 0.5. In all cases, each point represents a target, and 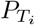 is obtained from averaging over 1000 independent realizations in the same environment. Ten different landscapes are shown. All the simulations were performed with *N* = 200, *γ*_*max*_ = 0.9, and *β* = 1.

**Fig 5.**
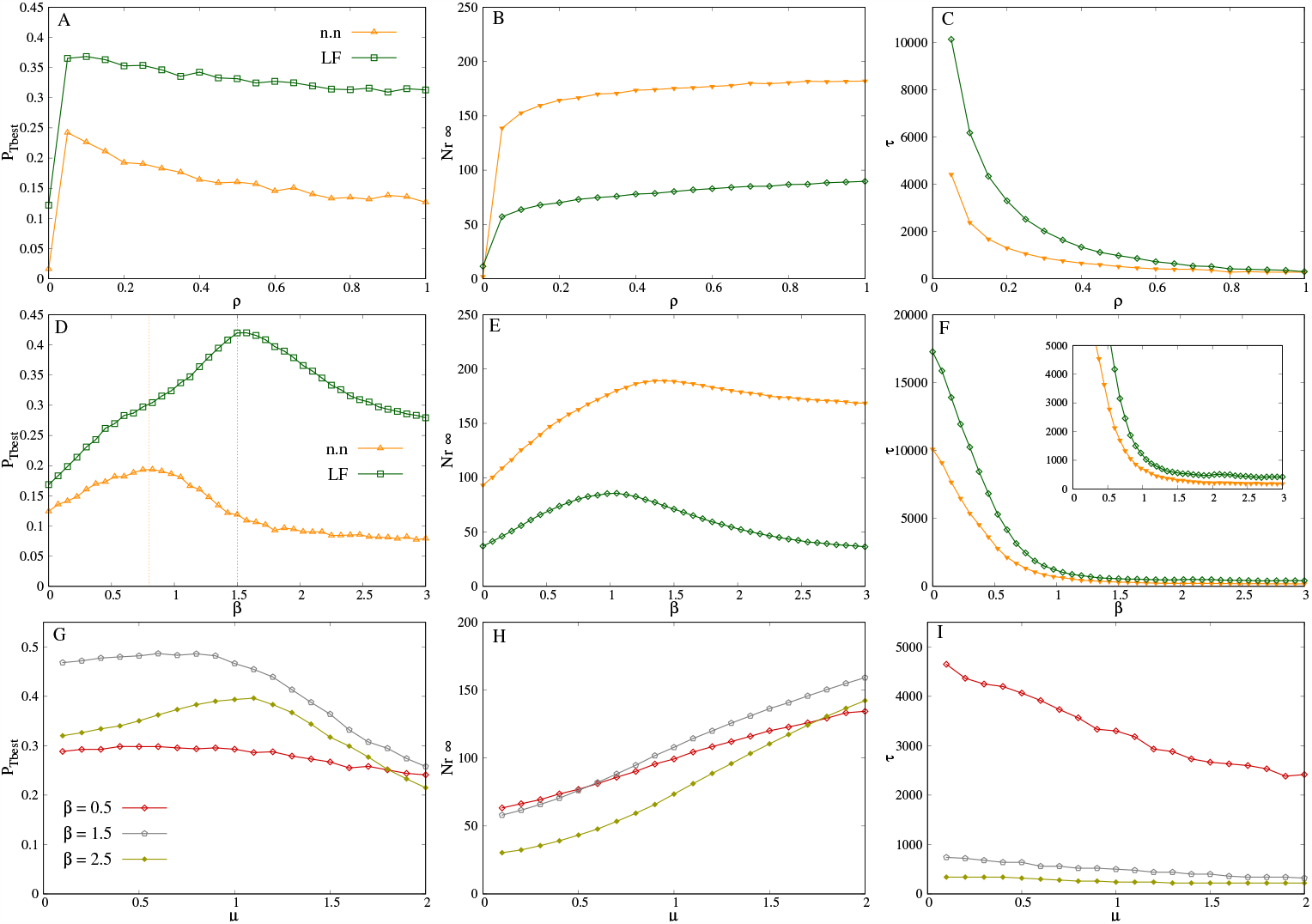
Memory decay case. Occupation probability of the best target 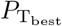 (left column), asymptotic cohesion 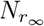 (center column), and typical learning time *τ* (right column), as function of: (A-C) the rate of information transfer *ρ* (with *µ* = 0.5 and *β* = 1); (D-F) the memory decay exponent *β* (with *ρ* = 0.5 and *µ* = 0.5); and (G-I) the Lévy flight index *µ*, for *ρ* = 0.5 and *β* = 0.5 (red line), *β* = 1.5 (grey line) and *β* = 2.5 (green line). In all cases, the memory rate was fixed to *q* = 0.25. In panel D, the values of the optimal memory decay exponent *β*^*∗*^ are indicated in vertical dashed lines for n.n. (yellow) and LF (green) swarms. The other parameters are those of Fig 2. All the curves are averages over ten different landscape configurations.

To discuss the properties of n.n. and LF swarms with memory decay, we computed 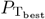, 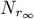, and *τ* as a function of the communication rate *ρ*, the memory decay index *β*, and the Lévy index *µ*, respectively.

#### Behavior as a function of the information transfer rate ρ

For both n.n. and LF swarms, when *β* is fixed to 1, the probability 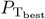 is approximately doubled compared to the case *β* = 0 (Fig 5A vs. Fig 3A). It also remains remarkably constant as a function of *ρ* (as far as *ρ >* 0), and slightly *decreases* with increasing the frequency of interactions. The curves of the cohesion 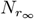 in Fig 5B reach plateau values at large *ρ* that are nearly twice the ones of Fig 3B. One of the most spectacular effects of memory decay, though, is the decrease of the learning time *τ* by a factor ranging between 5 (at *ρ* = 0.1) and 12 (at *ρ* = 0.4), compared to the perfect memory case (see the scale of the y-axis of Fig 5C vs. Fig 3C). The results presented in Fig 5A, B and C clearly show that the quantities 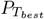, 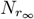, and *τ* are impacted by the memory decay exponent *β*.

#### Behavior varying the memory decay exponent β

To further investigate the behavior of the system when varying the exponent *β* of the memory kernel (see Eq. 2), we computed 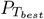, 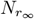, and *τ* as functions of *β* for fixed values of *ρ* = 0.5, *q* = 0.25, and *µ* = 0.5. Fig 3D shows the occupation probability of the best target 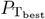 as a function of *β* for n.n. (yellow curve) and LF (green curve). In both dynamics, intermediate values of memory decay improve localization and, notably, 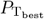reaches a maximum for a particular value of *β* (*β*^*∗*^ *≈* 0.8 for n.n and *β*^*∗*^ *≈* 1.5 for LF). The improvement in localization properties brought by memory decay is more significant for LF swarms, as the maximum probability 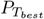 is always considerably larger for Lévy flights than for nearest neighbors. In the former case, the occupation probability of the best site at optimal, *β* = *β*^*∗*^, is 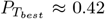, which is larger than the value with *β* = 0 by a factor of 2.5. In both dynamics (n.n and LF), a high memory capacity (*β ≈* 0) actually causes the group to reinforce those sites visited at the beginning of the search, which are not necessarily the most attractive sites. On the other hand, at low memory capacity (*β >* 3) the motion of individuals becomes similar to memory-less random walkers.

Similarly to 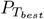, the cohesion described by the stationary number of neighbors 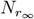, reaches maximum values close to *β≈* 1 for n.n. walkers, and *β≈* 1.5 for LF walkers. (See Fig 5E). As can be seen from this figure, the n.n. swarms are again more cohesive than LF ones. Note that the maximal cohesion is not reached at the exact same value *β*^*∗*^ reported in Fig 5D. Nevertheless, high levels of aggregation are associated with an intensive exploitation of the best resource site.

As mentioned before, the learning time *τ* drastically decreases with *β*, until a plateau is reached at about *β ≃* 1, see Fig 5F. When individuals have an infinite memory capacity (*β* = 0), the group takes a considerable longer time to cluster around the best resource sites.

In summary, by choosing *β≃* 1 for n.n. swarms and *β ≃*1.5 for LF swarms, a good compromise is achieved for a quick convergence, a selective exploitation and a high group cohesion. In addition to the power-law memory kernel analyzed above, we also performed simulations with an exponential kernel 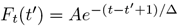 with no qualitative difference in the results (see Fig A in S1 Text).

#### Behavior as a function of the Lévy flight index µ (length of the steps)

To address the effects of the long exploratory random jumps on the group properties, we computed the three quantities above, 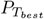, 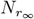, and *τ*, as functions of the Lévy index *µ* (that determines the length of the steps), for different values of *β* (which determines the memory kernel function). In Fig 5G, with a relatively slow memory decay (*β* = 0.5), the occupation probability 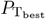 stays almost constant (around 0.3) as *µ* varies. With a more pronounced forgetting (*β* = 1.5 and *β* = 2.5), 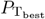 becomes non-monotonic with *µ* and reaches a maximum value around *µ* = 1. On the other hand, one can observe that swarm aggregation increases as the individuals perform shorter steps (have a larger *µ*) for all the values of *β* considered (Fig 5H). The cohesion of a swarm with slowly diffusing individuals is indeed not disrupted by large steps, which are more frequent in the highly super-diffusive regime (small *µ*). Finally, Fig 5I shows that the learning time *τ* slightly decreases with *µ*. In this last example, the swarm with the slower forgetting capacity (*β* = 0.5, red line) has a learning time considerably longer than the other two.

## DISCUSSION

Groups of social agents are often more effective in solving complex tasks than isolated individuals, not only in ecology [52] but also in the area of artificial intelligence [57] or in the context of the discovery of innovations, where social connections play an important role [58]. Individual experience and the transfer of valuable information that can be exploited by others are essential in the emergence of collective learning and novelties in human societies [58]. In the present study, individuals discover a foraging site not visited previously as a result of their random exploration of the environment. They also store information about previously visited sites, and, depending on their memory capacity, retrieve this information at a constant rate to take memory-based decisions.

With these elements, we developed a model of interacting agents with memory and observed the emergence of collective states of learning. We analyzed different individual exploratory behaviors and memory features, used by the foragers to visit resource patches according to their attractiveness. As a consequence of pairwise interactions, in all cases the whole group gathered very selectively around the best resource site among the many sites available in the environment, except when individuals made little use of their memory. The reward obtained by the individuals of the group is much higher than what they would obtain on their own without sharing information. We have explored the effects on learning of the use of random walks or Lévy flights, as well as the effects of a perfect memory capacity or a memory that decays over time as a power law.

In the perfect memory case, the cohesion of n.n swarms converges faster and to higher values than in LF swarms (larger 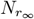 and smaller *τ*). However, LF groups localize more strongly on the most attractive target (larger 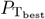), meaning that the group takes advantage of the valuable information collected by individuals, which at the same time is facilitated by their long exploratory steps.

When memory decays with time, we found qualitatively similar results. Nevertheless, some progressive forgetting considerably reduces the time needed by the group to find and gather around the best resource sites. Concomitantly, individuals exploit these sites more intensively and are more cohesive. Therefore, a decay of memory allows the group to learn faster and better. In particular, an intermediate value of the decay exponent maximizes resource exploitation. If memory decays too fast, collective learning degrades and the group becomes similar to a swarm of interacting but memory-less random walks. Although n.n. swarms converge even faster than LF swarms to the steady state, they localize less effectively. Thus, LF movements with a proper amount of memory decay seem to represent an advantageous individual strategy for maximising the foraging success of the group (indicated by a combination of high 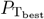 and small *τ*), as a result of efficient collective learning. This is in line with previous results on the benefits of a limited memory [34].

The ability to transfer information is essential to the emergence of collective learning. Surprisingly, the afore-mentioned efficient strategies require a fairly small amount of interactions among foragers. In comparison, stronger interactions (higher *ρ*) are needed to reach high levels of collective learning in the perfect memory case, see Fig 3A, 3B and 3C vs. Fig 5A, 5B and 5C.

The outcome of collective actions is not always rewarding and can be costly instead. Inspired by the homing behavior of pigeons, a model of two coupled neural networks aiming to learn a correct navigation angle showed that the performance of solo learners was actually better than the collective performance of democratic pairs of learners [59]. However, when pair members contributed less equally to decision making, learning was faster and final performance higher [59]. In the present model, collective performance might be a consequence of the complexity of the foraging environment and of the large number of individuals, who can explore different regions of space in parallel. However, the (slight) decrease of performance with the strength of interactions, observed in Fig 5A, while the cohesion increases in Fig 5B, is worth being noticed. This sub-optimal patch exploitation due to an excess of interactions could be caused by the presence of conflicting information in the social network about the location of the best resource sites.

A wealth of theoretical studies have discussed the optimality of random Lévy walks for encountering resources in unknown environments [9, 60–63]. Here we have shown that multiple scale movements can also be useful for learning purposes, when dispersed information about resources needs to be collected during a memory-based process. In a model similar to ours, where individuals performed random exploratory Lévy walks at constant speed (instead of jumping from one site to another in one time-step like here), a large fraction of the group could also aggregate around a salient resource site of the environment [53]. Aggregation even reached a maximum for a particular value of *µ*, which decreased as the use of memory was more frequent. Here we find that increasing *µ* favours cohesion, at the expanse of less exploitation of the best site.

In social species such as spider monkeys, the individuals have mental maps to navigate their environment and also exhibit movements characterized by a Lévy index of *µ ∼* 1 [12, 55]. Spider monkeys travel in small subgroups of varying composition and size across their home range and can communicate over large distances through vocalizations [64]. Information transfer processes and their effects on collective dynamics have also been measured in this species [65]. By monitoring visits to certain food patches, knowledgeable and naive individuals can be identified, and it has been demonstrated that the arrival of naive individuals to novel food patches is accelerated by social information [65]. Our model exhibits a similar phenomenon, as illustrated by Fig 2C and 2D regarding the number of individuals that have visited a particular site at a given time, in the presence or absence of interactions.

Information transfer in the context of collective foraging can be active, as when overt signals are used specifically to recruit others to feeding areas, or passive, as when individuals simply follow others to known areas [66]. The latter is the simplest mechanism by which information could be transferred, and would imply that information would be distributed more equally among individuals, while active signalling could involve unequal behavior on the part of signallers and the responses by recipients. Dominant individuals, for example, could signal freely as they would not be displaced from small patches, while recipients could decide to join or not depending on their relationship to the signaller [67]. In our model, we do not assume any particular mode of information transfer, but it would be compatible with an equally distributed way of information transfer. In the future, it would be interesting to study how collective performance changes when other types of information transfer are considered.

It is well accepted that forgetting improves reinforced learning because unimportant information disappears, making room for more valuable input [36–38, 68]. Here, we have allowed memory to decay slowly in time as a power-law instead of exponentially. The classical Ebbinghaus’ “forgetting curve” obtained from experiments conducted on human subjects is well fitted by a double power-law or a simple power-law function with a small exponent (in the interval 0.1 *−* 0.2), rather than by an exponential or a double exponential [69]. The collective decay of public memories in human societies can be described as an exponential at short time and a power-law with *β* = 0.3 at large time [70]. In the context of movement ecology, the ranging patterns of bisons are well fitted by power-law memory kernel with *β* = 1 [1]. Similar data of Elk movements were successfully fitted by a slower-than-exponential decay [27]. With the model proposed here, we have shown that memory decay is an important driver of a fast and efficient collective learning in static but complex environments. When assuming a different memory kernel, with exponential decay, we found qualitatively similar results (see Fig A in S1 Text).

In its present form, our computational model considers resource patches that are always available and ignores competition among foragers. However, even the best food patch may not be sufficiently large for being occupied by a large number of individuals at the same time. In addition, patch depletion by some foragers may affect the attractiveness perceived by others during future visits, unless the patches refresh very rapidly. These competition effects should favour the occupation of other good but sub-optimal patches with a larger probability and cause a decrease in cohesion by splitting the group into several subgroups, akin to fission-fusion societies [71]. We nevertheless expect that medium-sized groups should still choose patches very selectively according to the attractiveness, even in the presence of competition.

On the other hand, travel costs play a crucial role in shaping an individual’s decision-making process within the context of animal foraging. When an animal lacks comprehensive information about the attributes of the destination, the inclination to undertake an extended journey could be diminished. We modified the primary model, incorporating a travel cost correlating with the distance between the individual’s current position and the target destination. The outcomes of this adjustment are illustrated in Fig B in S1 Text of the Supplementary Information. Notably, the results reveal no significant alterations in the collective learning performance across the swarm, thus highlighting the robustness of the model’s findings.

Even though our work represents a step toward the understanding of some mechanisms responsible for efficient collective learning in ecology, other aspects remain elusive. The effects of the number of foragers on the learning time and performance are little understood. We speculate that the foraging efficiency of a group increases rapidly with the number of individuals, as the search for good resource sites is faster in principle with foragers dispersed over a whole landscape. However the fact that too many interactions seem to affect learning would suggest that beyond a certain size, the group could find conflicting information and strive to make optimal decisions. Future studies could also focus on how the foragers of our model adapt to environments where resources are clumped into patches or that are ephemeral. In changing environments, the decay of memory might be particularly beneficial.

## ACKNOWLEDGEMENTS

AFC thanks A. Aldana for fruitful discussions. We thank Carlos E. López Natarén for technical computer support.

## SUPPORTING INFORMATION

**S1 Text**. Extensions of the model: expoential memory decay and travel costs.

**Fig A in S1 Text. Exponential memory decay**. Simulations with *N* = 200 walkers in an environment with 200 *×* 200 lattice sites, resource density *δ* = 0.0025 (*M* = 100 resource sites), *γ*_*max*_ = 0.9 and *t* = 10^5^. (a) Occupation probability of the best target, 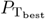, (b) Asymptotic cohesion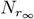, and (c) Learning time *τ* for n.n. (green) and LF dynamics (yellow), as a function of the memory time ∆. The other parameters are *q* = 0.25, *ρ* = 0.5 and *µ* = 0.5. All the curves are averages over one thousand different walks in one landscape configuration.

**Fig B in S1 Text. Cost for long travels**. (a) Occupation probability of the best target, 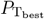, (b) Asymptotic cohesion 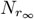, and (c) Learning time *τ* as function of the rate of memory use *q*, with *ρ* = 0.5. The swarm performs a LF dynamics with *µ* = 0.5. The other parameters are those of Fig A. All the curves are averages over one thousand different walks and only one landscape configuration.

## Notes

### Competing Interest Statement

The authors have declared no competing interest.

### Summary of Updates

Extra subsections were added in order to a better understanding of Figs. 3 and 5. Also, an SI section is now presented to address other important aspects of the agent-based model.

