## Supplementary Information for "Lévy movements and a slowly decaying memory allow efficient collective learning in groups of interacting foragers"

October 14, 2023

### 1 Exponential memory kernel

Fig A shows the result obtained from averaging over 1,000 independent swarms exploring a same environment, where we have considered that each individual has a memory kernel following an exponential decay, *i.e.*, $F_t(t') = Ae^{-(t-t'+1)/\Delta}$, with $\Delta$ a characteristic decay time and $A$ a normalization factor. Noticeably, the results are qualitatively similar to those obtained with the other two memory kernels presented in the main text. Lévy Flight (LF) swarms are roughly three times as efficient as nearest neighbors (n.n.) swarms, for any value of $\Delta$, in the task of finding the best resource site in the environment (Fig AA. As before, n.n. swarms reach a better (about twice larger) asymptotic cohesion $N_{r_\infty}$ (Fig. AB and have a slightly shorter learning time $\tau$ (Fig. AC than LF swarms. The plateaus in curves A and B as $\Delta \to \infty$ correspond to the infinite memory case.

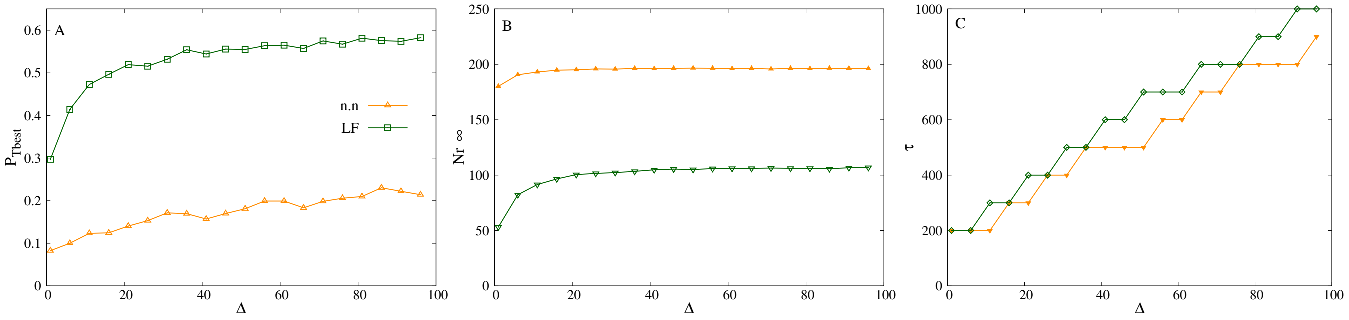


Fig A : *Exponential memory decay.* Simulations with $N = 200$ walkers in an environment with $200 \times 200$ lattice sites, resource density $\delta = 0.0025$ ($M = 100$ resource sites), $\gamma_{max} = 0.9$ and $t = 10^5$. A) Occupation probability of the best target, $P_{T_{best}}$, B) asymptotic cohesion $N_{r_\infty}$, and C) learning time $\tau$ for n.n. (green) and LF dynamics (yellow), as a function of the memory time $\Delta$. The other parameters are $q = 0.25$, $\rho = 0.5$ and $\mu = 0.5$. All the curves are averages over one thousand different walks in one landscape configuration.

### 2 Effects of travel costs

To analyze the possible impact of travel costs on the results of the main text, we have implemented an additional rule that discourage long movement steps. We assume that, in the exploration phase, an individual chooses to go to a site $m$ from a site $n$ with probability $1/d(n,m)$, where $d(n,m)$ is the Euclidian distance between the two sites. If the individual chooses not to perform this step, it picks up a new site until the step actually takes place. Therefore, the probability of traveling to a new site decay linearly with the distance, and LF steps are more penalized than n.n. steps.

Fig B shows the main swarm performance metrics studied in the main text as a function of the memory use parameter $q$. Here we consider a LF swarm with infinite memory. The blue curves correspond to a swarm with travel costs, as presented in the main text, whereas the pink curves correspond to a swarm that incorporates travel costs. The two cases exhibit the same qualitative behaviors. Notably, $P_{\mathrm{T_{best}}}$ is even slightly larger with travel costs, although the learning time is longer. Overall, as far as collective learning is concerned, the swarm is not significantly affected by travel cost penalization.

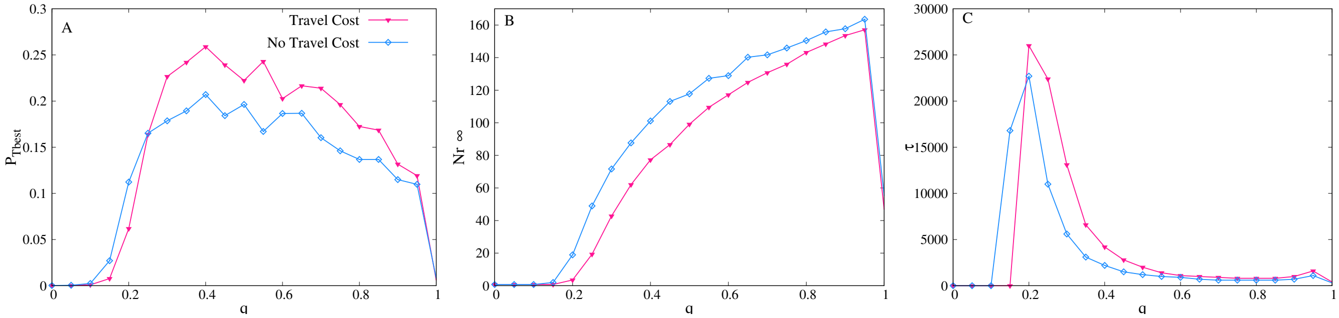


Fig B : *Cost for long travels.* A) Occupation probability of the best target, $P_{\mathrm{T_{best}}}$, B) asymptotic cohesion $N_{r_\infty}$, and C) learning time $\tau$ as function of the rate of memory use $q$, with $\rho = 0.5$. The swarm performs a LF dynamics with $\mu = 0.5$. The other parameters are those of Fig. A. All the curves are averages over one thousand different walks and only one landscape configuration.
